# Assessment of the petroleum hydrocarbon biodegradation potential of the sediment microbial community from an urban fringing tidal marsh of Northern New England

**DOI:** 10.1101/279760

**Authors:** Sinéad M. Ní Chadhain, Jarett L. Miller, John P. Dustin, Jeff P. Trethewey, Stephen H. Jones, Loren A. Launen

## Abstract

We assessed the impact of dodecane, *n-* hexane and gasoline on the microbial diversity of chronically polluted fringing tidal marsh sediment from the Great Bay Estuary of New Hampshire. Dilution cultures containing saturated alkane concentrations were sampled at zero, one and 10 days, and *alkB* and *cyp153A1* alkane hydroxylase gene libraries and 16S rRNA sequences were analyzed. The initial sediment had the most diverse alkane hydroxylase sequences and phylogenetic composition whereas treated sediments became less functionally and phylogenetically diverse with alkane substrates apparently enriching a few dominant taxa. All 1-and 10-day samples were dominated by *Pseudomonas-* type alkane hydroxylase sequences except in dodecane treatments where primarily *Rhodococcus-*-type alkane hydroxylases were detected. 16S rRNA profiling revealed that the Gammaproteobacteria, particularly *Pseudomonas*, dominated all one day samples, especially the *n-* hexane and gasoline treatments (63.2 and 47.2% respectively) and the 10-day *n-* hexane treatment (which contained 60.8% *Pseudomonas* and 18.6%*Marinobacter).* In contrast, the 10 days of dodecane treatment enriched for Actinobacteria (26.2% *Rhodococcus* and 32.4%*Mycobacterium)* and gasoline treatment enriched for Firmicutes (29.7%; mainly *Bacillus, Lysinibacillus* and *Rumelibacillus).* Our data indicate that fringing tidal marshes contain microbial communities with alkane-degrading abilities similar to larger meadow marshes, and support the hypothesis that alkane exposure reduces the functional and phylogenetic diversity of microbial communities in an alkane-specific manner. Further research to evaluate the ability of such fringing marsh communities to rebound to pre-pollutant diversity levels should be conducted to better assess the threat of petroleum to these habitats.

## Introduction

Salt marshes are frequently contaminated with petroleum through both large-scale releases, such as the Deepwater Horizon well blowout in the Gulf of Mexico in 2010 (Atlas et al., 2015; Natter et al., 2012; McGenity 2014), as well as chronic lower-level contamination from sources such as stormwater and small marine transportation related spills (McGenity 2014; Vieites et al., 2004). Microbial degradation of petroleum hydrocarbons released into marshes is the major means of removal, as evidenced by research demonstrating both the loss of petroleum, and the expansion of hydrocarbon-degrading taxa and associated degradative genes (Acosta-Gonzalez et al,. 2015; Atlas et al., 2015; Beazley et al., 2012; Kimes et al., 2014, Koo et al., 2015; Looper et al., 2013; Lu et al., 2012; Mahmoudi et al., 2013; Vega et al., 2009; Zhu et al., 2004). Petroleum bioremediation potential is related to the chemical composition of the polluting petroleum, the composition of the indigenous microbial communities present in the receiving environment, and factors such as marsh vegetation (Acosta-Gonzâlez et al., 2015; Atlas 1975; Atlas et al., 2015; Beazley et al., 2012). Petroleum contains hundreds of different hydrocarbons, including high quantities of *n-* alkanes (Speight 1998). *N-* alkanes, are, therefore, one of the major groups of contaminants that routinely intrude into salt marshes.

*N-* alkanes are some of the most biodegradable petroleum hydrocarbons, particularly under aerobic conditions where many bacteria, particularly α, β and γ-Proteobacteria, Actinomycetales and Firmicutes, can degrade a range of *n-* alkanes (Joye et al., 2016; Mason et al., 2012; Mukherjee et al., 2017; Rojo 2009; Van Beilen & Funhoff 2007; Wang et al., 2010). Some alkane-degrading bacteria are actually obligate or near-obligate alkanotrophs such *Alcanivorax borkumensis* (Sabirova et al., 2006). Aerobic degradation of medium chain-length alkanes (C5-C16), is initiated by terminal carbon hydroxylation (Van Beilen & Funhoff 2007), effected by two types of alkane hydroxylases: membrane-bound non-haem diiron alkB-type hydroxylases (Kok et al., 1989; Van Beilen et al., 1994), and soluble Class 1 cytochrome P450 Cyp153A1 (CYP *153A1)* hydroxylases (Asperger et al., 1981; Kubota et al., 2005; Maier et al., 2001). The *alkB* and CYP *153A1* genes have been characterized in isolates and environmental DNA and in the case of *alkB* are considered indicators of enhanced microbial hydrocarbon degradation activity (Lu et al., 2012; Mukherjee et al., 2017; Van Beilen & Funhoff 2007, and others). Some bacteria possess only *alkB*, others only CYP *153A1*, and some have multiple alkane hydroxylase systems that work in concert (Chen et al., 2017, Liu et al., 2011; Nie et al., 2014; Schneiker et al., 2006; Van Beilen and Funhoff 2007; Wang et al., 2010a). In addition to *alkB* and *CYP 153A1* novel alkane hydroxylases continue to be discovered (Guibert et al., 2016 and others).

The majority of studies on petroleum degradation by salt marsh microbial communities have focussed on large scale release events due to shipping accidents or blowouts and large salt marsh meadow systems such as those found in the Gulf of Mexico area impacted by the 2010 Deepwater Horizon blowout (McGenity 2014). In New Hampshire, small estuarine marshes referred to as fringing marshes comprise a significant amount of the total salt marsh habitat (Morgan et al., 2009, PREP 2013) which have important ecosystem functions and values (Morgan et al., 2009). These fringing marshes are different from large meadow marshes, possessing lower levels of organic carbon and plant density (Morgan et al.,2009) and lower levels of denitrification enzyme activity (Wigand et al., 2004). Although fringing marshes are the first location impacted by petroleum influx, which is considered one of the major threats to such marshes in New Hampshire (NHDES, 2004), little is known about the petroleum biodegradation potential of these marshes. In this study we aimed to assess the catabolic and phylogenetic diversity of an alkane-degrading fringing marsh microbial community from a brackish chronically petroleum-impacted fringing marsh site on the Cocheco River of the Great Bay Estuary of New Hampshire (Watts et al.,2006). Furthermore we aimed to evaluate the response of the community to different sources of *n-* alkanes (gasoline, *n-* hexane and dodecane). We hypothesized that given the chronic nature of petroleum input in the area, the indigenous microbial community would contain diverse alkane hydroxylase genes, and that exposure to different *n-* alkanes would select for different suites of alkane hydroxylase genes, reducing gene-specific and overall phylogenetic diversity through selection of degradative members of the community. To test this, we treated dilution cultures to each source of *n-* alkanes for 10 days, analyzing *alkB* and *CYP 153A1* alkane gene fragments from clone libraries, and 16S rRNA genes to assess changes in the functional and taxonomic structure of the community respectively.

## Materials and Methods

### Sediment sampling and dilution cultures

Sediment samples were collected from a tidal marsh vegetated with *Spartina alterniflora*, located along the Cocheco River in Dover, NH (43°11’5L36”N 70°52’02.81”W) on 11 May 2011 (after spring plant emergence) at low tide. The Cocheco River is listed as impaired (NHDES 2015) due to the chronic input of petroleum hydrocarbon contamination from shipping and other sources (Magnusson et al.,2012) as well as historical and current point and non-point sources including waste water treatment plants, landfill leachate (such as the Superfund listed Tolend Road landfill in Dover NH: EPA) and urban stormwater. Because the level of impervious surfaces is increasing (PREP 2013) stormwater-carried contaminants, such as petroleum hydrocarbons (Brown et al.,2006; Makepeace et al.,1995), routinely enter the river and its tidal marsh sediment. Several sediment samples were collected from the top 15 cm of a 3 m x 3 m area containing areas vegetated and un-vegetated areas and composited into a sterile 1 gallon container. Large plant material, rocks or other visible detritus was removed by hand and the sample was homogenized by passing through a Food Mill (RSVP International Inc, Seattle, Washington) using two different filtering disks with pore size 0.5 cm and 0.2 cm. Dilution cultures were established by adding 1 g (wet weight) sediment into 50 mL modified Minimal Salts Broth (Launen et al., 2008) in 250 mL Erlenmeyer flasks. Dilution cultures received (separately) 1.5 mL gasoline (G) or *n-* hexane (H) in the vapor phase, dodecane (D) neat to saturating concentrations or no alkanes (no-alkane control). All *n-* alkanes were maintained to ensure saturating concentrations throughout the 10 days of exposure. Room temperature cultures were shaken at 100 rpm to ensure aerobic conditions and were sampled at time 0 (initial, T0), and after 1 (T1) and 10 (T10) days for genomic DNA extraction and subsequent processing.

### DNA extractions and clone library preparation and sequencing

Genomic DNA was extracted from 4 mL of dilution cultures using the PowerSoil DNA Isolation Kit (Mo Bio Laboratories, Carlsbad, CA). PCR was conducted using primers that targeted the *alkB* alkane hydroxylase (Kloos et al.,2006; Van Beilen et al.,2006; Wang et al.,2010b). The forward primer of Kloos et al.,was modified by addition of an R on the 3’ end. PCR reactions were conducted using 1μM of forward and reverse primers, 2X Green GoTaq Reaction Buffer (Promega, Madison, WI), 1 μL template DNA (ca. 10 − 50 ng) and nuclease-free water (to 20 μL). The amplification cycle consisted of an initial denaturing step of 95°C for 5 minutes, followed by 35 cycles of 95°C for 30 seconds, 55°C for 30 seconds and 72°C for 30 seconds, and a final elongation of 72°C for 5.5 minutes.

Hydroxylase gene fragments were cloned into pCR4-TOPO vectors (Life Technologies, Carlsbad, CA). Plasmid DNA was isolated using the PureYield Plasmid Miniprep System (Promega). Sequencing was conducted by the University of Washington High Throughput Genomics Center using either the M13 forward or reverse primers provided by the sequencing facility.

### Phylogenetic analysis of clone library sequences

Primer sequences were removed from nucleotide sequences using Geneious v. 7.1.5 (created by Biomatters). Sequences were aligned with a ClustalW alignment (in Geneious) and Phylip v. 3.695 (Felsenstein 2005). An alignment-associated DNA distance matrix was input into the program MOTHUR v. 1.33.1 (Schloss & Handelsman 2009), that was used to cluster sequences into operational taxonomic units (OTUs) based on the nearest neighbor algorithm and a 95% (nucleotide) sequence identity. MOTHUR was also used to construct rarefaction curves for each library, and to calculate estimates of nonparametric richness (Chao1 and Shannon-Weaver diversity indices) (Chao 1984; Chao and Lee 1992; Pielou 1977). Phylogenetic trees were constructed from the ClustalW alignment of the derived amino acid sequences by neighbor-joining analysis with 1000 bootstrap replicates using MEGA 5.0 (Tamura et al.,2011).

### Nucleotide sequence accession numbers

The nucleotide sequences reported in this study were deposited in the GenBank database with the following accession numbers KT280353 to KT280397 (selected representatives of each *alkB* OTUs) and KT305744 to KT305774 (selected representatives of each *CYP 153A1* OTUs).

### Quantification of relative *alkB, CYP 153A1* and 16S rRNA gene abundance in dilution cultures

The dominant *alkB* and *CYP 153A1* OTUs from the clone libraries were further analyzed using real-time PCR. QPCR primers (Supplemental Table 1) were designed from the nucleotide sequences of clones from within the selected OTUs using Primer Express (V3.0.1, Applied Biosystems). The abundance of selected *alkB* and *CYP 153A1* OTU sequences was determined relative to the abundance of bacterial 16S rRNA genes. Standard curves were generated using two representative plasmid clones from each OTU. Plasmid DNA was linearized with the *NotI* restriction enzyme and quantified using a Nanodrop 2000 (Thermo Orion). Standard curves for bacterial 16S rRNA genes were generated using *Escherichia coli* JM109 genomic DNA. QPCR reactions were performed in 96-well Fast Optical MicroAmp reaction plates (Life Technologies) where each reaction had a total volume of 12.5 μl: 1 μl of environmental sample or plasmid DNA solution, 6.25 μl SYBR Green PCR Master Mix (Applied Biosystems), 4.75 μl water and 0.25 μl of each primer. Thermal cycling and quantification were performed using a StepOne Plus Real-Time PCR System (Applied Biosystems) instrument programmed as recommended by the manufacturer (95°C for 10 min followed by 40 cycles of denaturation at 95°C for 30 s, primer annealing at the primer-specific annealing temperature (Table 3) for 1 min, and polymerase extension at 72°C for 1 min. The final primer concentration was 0.2 mM. Amplification specificity was achieved for each primer set, as determined by the presence of a single peak in each post-amplification dissociation curve.

### Bacterial community sequencing

PCR amplification of the 16S rRNA hypervariable region V6 was performed with a pool of degenerate forward and reverse primers targeting bacteria as described by Huber at al., (2007). The 5’-ends of the forward primers were fused with the Ion Torrent A-Adaptor plus a MID key sequence (barcode), while the reverse primers were fused with the truncated P1-adapter sequence (trP1). The primers were diluted in molecular grade laboratory water and pooled in equimolar concentrations for each barcode set. Template from each treatment was amplified with a different barcode.

For amplicon library preparation purified genomic DNA from each treatment was amplified in 20 μL reactions as follows: 1 μL of DNA (10-50 μg/mL concentration), 10 μL 1X Phusion PCR master mix (Thermo, Pittsburgh, PA), and 2 μL of 0.5 μM forward and reverse primers. The PCR conditions were: 94°C for 3 min, followed by 30 cycles of 94°C for 15 s, 58°C for 15 s, 68°C for 10 s, and a final elongation step of 68°C for 30 s. Triplicate amplifications were performed for each treatment. PCR products were run on 2% 1X TBE gels. Amplicons of the correct size were excised, pooled, and purified with the Promega SV Gel and PCR Clean-Up kit (Promega, Madison, WI). Amplicon concentration was estimated with the Qubit 2.0 instrument using the Qubit dsDNA HS assay (Life Technologies). The bar coded amplicons were combined and diluted to 2.8x10^s^ DNA/μL molecules per micro liter prior to template preparation.

Sequencing template was prepared using the Ion OneTouch 200 Template Kit with the corresponding protocol (Pub. Part no. 4474396 Rev. B, 20 Feb 2012). Sequencing of the amplicon libraries were then carried out using the Ion Torrent Personal Genome Machine (PGM) system using the Ion Sequencing 200 kit (from Life Technologies) and the corresponding protocol (Pub Part Number 4474596 Rev. C, 10 Oct 2012) on Ion Torrent 314 chips.

### Bacterial community sequence analysis

Raw sequencing reads were processed using the Quantitative Insight Into Microbial Ecology (QIIME) open source software package and the Virtual Box QIIME package (Caporaso et al., 2010a). Raw sequences were filtered and screened for different quality criteria. Low quality reads were removed as follows: i) all reads <70 b.p. were removed; ii) all reads with quality scores of lower than 20 were removed; iii) no ambiguous reads were allowed; and iv) reads with homopolymers of 4 b.p. or less were removed to reduce sequencing error. Chimeras rarely form in the V6 regions of the sequence due to its small size (approximately 70-200 b.p. sequence lengths) so the data were not screened for chimeras.

Subsequently, de novo OTUs were clustered at a 97% identity threshold and aligned to the Greengenes database (DeSantis et al 2006) using PyNAST (Caporaso et al., 2010b). Alpha diversity was calculated using the QIIME pipeline. The default metrics for the QIIME pipeline alpha diversity measurement were edited to include the Shannon diversity index and phylogenetic distance, in addition to Chao1 and observed species. The minimum number of sequences used for depth of coverage in alpha rarefaction was set to 5000 for even sampling. Beta diversity metrics were also generated using QIIME. 5000 sequences were subsampled from each dilution sample for calculation of beta diversity using both weighted and unweighted unifrac (Lozupone & Knight 2005) analyses. Principal Coordinate Analysis (PCoA) (Vazquez-Baeza et al., 2013) and hierarchical clustering (UPGMA) (Caporaso et al., 2010a) were utilized to visualize the data structure and differences between the samples. Jackknife replicates (100 replicates) were used to estimate the uncertainty in both the PCoA and hierarchical clustering plots.

## Results

### Analysis of alkane hydroxylase clone libraries

#### *alkB* clone library

*AlkB* alkane hydroxylase gene fragment clone libraries were prepared from genomic DNA extracted from alkane-exposed and unamended cultures over 10 days. The distribution of OTUs, their diversity, and matches to known sequences as determined by Blastx are shown in Table 1 and 2, and Fig. 1A and 2. Dilution culture alone reduced diversity of *alkB* gene fragments according to the Shannon-Weaver diversity index and Chao1 estimates (Table 1) in the first day of dilution culture, however, both measures increased by the T10 to levels greater than the initial T0 sediment indicating that diversity rebounded in the dilution culture. In contrast, all alkane treatments reduced *alkB* gene fragment diversity to levels less than those found in the no alkane control sample after 1 and 10 days of continuous alkane exposure.

**Table 1.** Diversity analysis of functional gene clone libraries. OTUs were clustered at the 95% identity level using the furthest neighbor in Mothur version 1.33.3

| Library | N | N | H' | Chao1 |
| --- | --- | --- | --- | --- |
| *alkB* |  |  |  |  |
| TO NA | 25 | 14 | 2.44 | 23 |
| T1 NA | 14 | 9 | 2.04 | 14 |
| T1 H | 29 | 8 | 1.75 | 8 |
| T1 D | 33 | 9 | 1.81 | 19 |
| T1 G | 31 | 4 | 0.65 | 5 |
| T10 NA | 26 | 15 | 2.46 | 42.5 |
| T10 H | 24 | 2 | 0.56 | 2 |
| T10 D | 19 | 3 | 0.66 | 3 |
| T10 G | 21 | 5 | 0.87 | 6.5 |
| *CYP 153A1* |  |  |  |  |
| TO NA | 13 | 13 | 2.56 | 91 |
| T1 NA | 3 | 3 | 1.1 | 1 |
| T1 D | 21 | 1 | 0 | 6 |
| T10 NA | 31 | 10 | 1.85 | 13.3 |
| T10 D | 1 | 5 | 1.13 | 5.5 |

<sup>a</sup>N, number of clones sequenced from each library; n, number of OTUs observed; H', Shannon-Weaver diversity index, Chao1 nonparametric richness estimate. Numbers in parentheses represent 95% confidence intervals; ND, not determined.

**Table 2.** Best BlastX matches (by identity) in the GenBank database for the dominant alkane hydroxylase clone OTUs. Where best match has no taxonomic information the first match with taxonomic information is also included.

| OTU # \|<br># of sequences in<br>OTU<sup>1</sup> | Gene ID | Name | Identity<br>(%) | e-value |
| --- | --- | --- | --- | --- |
| *alkB* |  |  |  |  |
| OTU1\|84 | WP_017846506.1 | *Pseudomonas veronii* | 100 | 3E-114 |
| OTU2\|16 | WP_026651424.1 | *Pseudomonas aeruginosa* | 81 | 3E-75 |
| OTU3\|15 | WP_026651424.1 | *Pseudomonas aeruginosa* | 84 | 3E-87 |
| OTU4\|15 | ABB90689.1 | uncultured bacterium | 99 | 8E-116 |
|  | AID55567.1 | *Rhodococcus* sp. RP-11 | 99 | 3E-115 |
| OTU5\|12 | CDI44588.1 | *Pseudomonas lini* | 96 | 6E-113 |
| OTU6\|7 | WP_038843879.1 | *Pseudomonas fluorescens* | 99 | 2E-113 |
| OTU7\|6 | ACT91140.1 | uncultured bacterium | 100 | 1E-119 |
|  | ADG26619.1 | uncultured *Rhizobiales* bacterium | 100 | 2E-119 |
| OTU8\|5 | AGQ20900.1 | uncultured bacterium | 80 | 9E-91 |
|  | ACJ22711.1 | *Alcanivorax* sp. S15-9 | 67 | 3E -80 |
| OTU9\|5 | ACJ22725.1 | *Kordiimonas gwangyangensis* | 81 | 5E-85 |
| OTU11\|4 | AGQ20978.1 | uncultured bacterium | 83 | 3E-95 |
|  | WP_027103373.1 | *Comamonadaceae* bacterium URHA0028 | 84 | 7E-94 |
| OTU13\|3 | AGQ20902.1 | uncultured bacterium | 82 | 3E -90 |
|  | WP_027103373.1 | *Comamonadaceae* bacterium URHA0028 | 81 | 7E -89 |
| OTU16\|2 | ABB96070.1 | uncultured bacterium | 86 | 1E -96 |
|  | WP_027103373.1 | *Comamonadaceae* bacterium URHA0028 | 85 | 1E-93 |
| OTU18\|2 | WP_030535917.1 | *Rhodococcus erythropolis* | 100 | 4E-112 |
| OTU19\|2 | BAG06232.1 | *Rhodococcus erythropolis* | 99 | 7E-117 |
| *CYP 153A1* |  |  |  |  |
| OTU1\|13 | CCO96780.1 | uncultured bacterium | 100 | 4E-64 |
|  | WP_032375608.1 | *Rhodococcus fascians* | 100 | 3E-63 |
| OTU2\|11 | ADF27209.1 | uncultured *Rhizobiales* bacterium | 93 | 2E-60 |
| OTU3\|8 | AKG58848.1 | uncultured bacterium | 88 | 2E-56 |
|  | ADO15009.1 | *Parvibaculum hydrocarboniclasticum* | 80 | 7E-51 |
| OTU4\|4 | ADF27251.1 | uncultured *Rhizobiales* bacterium | 93 | 2E-61 |
| OTU5\|3 | CCO96917.1 | uncultured bacterium | 98 | 4E-63 |
|  | ACP39704.1 | *Parvibaculum* sp. S18-4 | 97 | 7E-62 |
| OTU6\|2 | ADF27232.1 | uncultured *Rhizobiales* bacterium | 94 | 3E-48 |
| OTU7\|2 | CAH56117.1 | *Rhodococcus erythropolis* | 99 | 2E-64 |
| OTU16\|1 | CCO96881.1 | uncultured bacterium | 86 | 5E-55 |
|  | ADF27267.1 | uncultured *Rhizobiales* bacterium | 99 | 2E-53 |
| OTU27\|1 | ABI14020.1 | *Rhodoccus* sp. MOP100 | 87 | 1E-54 |
| OTU31\|1 | AKG58846.1 | uncultured bacterium | 78 | 1E-49 |
|  | ADO15009.1 | *Parvibaculum hydrocarboniclasticum* | 77 | 5E-49 |

<sup>1</sup>The number of matches to this OTU relative to all of the clone sequences from all libraries is shown. For example, there were 84 clone library sequences that comprised OTU1 when all libraries were considered.

**Figure 1.**
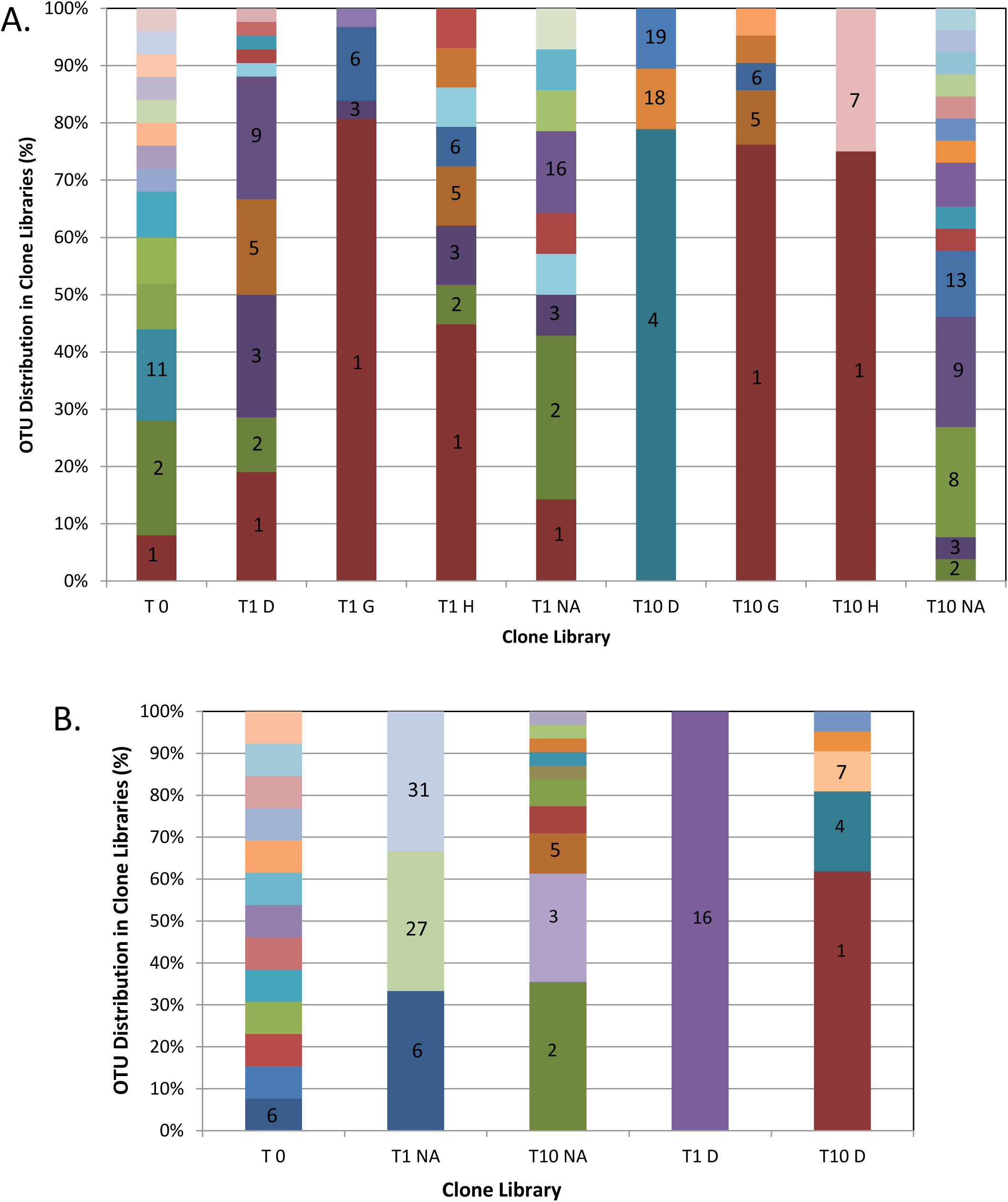
The distribution of *alkB* (A) and *P450* (B) OTUs in clone libraries derived from initial (T0), 1 day (T1) and 10 day (T10) samples exposed to dodecane (D), gasoline (G), *n*-hexane (H) or no alkanes (NA − control). Alignments, generation of distance matrices and clustering into OTUs at a similarity level of 95% was performed as described in the Methods. OTU numbering is included for OTUs that occurred in >10% of the total recovered for that library. See Table 2 for further information on OTU identity and Table 1 for the number of clone sequences in total for each library shown.

The taxonomic assignment of *alkB* sequences recovered from each sample library differed according to treatment (Table 2, Fig. 1A). The initial sediment (the T0 NA sample), yielded 25 *alkB* clone library sequences which contained 14 different OTUs, the majority of which were an 81% BlastX match to *alkB* from *Pseudomonas aeruginosa* (OTU2) or an uncultured *Commomodaceae* sequence (OTU11) (see Fig. 1A). After one day of dilution culture, the unamended control sample (T1 NA) yielded 14 clone library sequences, containing 9 OTUs, most of which matched OTU2 (P. *aeruginosa).* By 10 days of dilution culture alone (T10 NA) 26 *alkB* clone sequences were recovered, containing 15 OTUs, with the majority matching OTU8 (uncultured*/Alcanivorax* sequences as best match) and OTU9 *(Kordiimonas gwangyangensis*).

*AlkB* libraries prepared from *n-* hexane and gasoline amended samples yielded fewer OTUs overall and possessed reduced diversity measures relative to the unamended control libraries (Table 1). One day of exposure to both *n-* hexane (T1 H) and gasoline (T1 G) resulted in a selection for the *P.* veronii-like OTU1 (Fig. 1A, 14/29 and 25/31 of the clone sequences from these libraries respectively). This selection for *Pseudomonas-type alkB* sequences persisted into the 10 day samples where 18/24 (T10 H) and 16/21 (T10 G) were also sequences belonging to OTU1. Dodecane appeared to impact the *alkB* diversity differently than exposure to *n-* hexane or gasoline. While the one day dodecane-exposed sample was dominated by OTUs that match best to *Pseudomonas spp.* (Table 1 and Fig. 2A), by 10 days of dodecane exposure *alkB* sequences were predominantly OTU4 (15/19 sequences recovered), which is a 99% BlastX match to an *alkB* from *Rhodococcus* sp. RP-11 (Table 2, Fig. 1A). OTU4 was the major Gram positive type *alkB* identified in any *alkB* library prepared in this study. Overall, exposure to all of the alkanes tested resulted in differing profiles of *alkB* clone sequences than observed in the no alkane control samples, with *n-* hexane and gasoline treatment selecting for *alkBs* of *Pseudomonas-type* and dodecane treatment selecting initially for *Pseudomonas*-type sequences with a switch to *Rhodococcus*-type *alkB* sequences by day 10.

**Figure 2.**
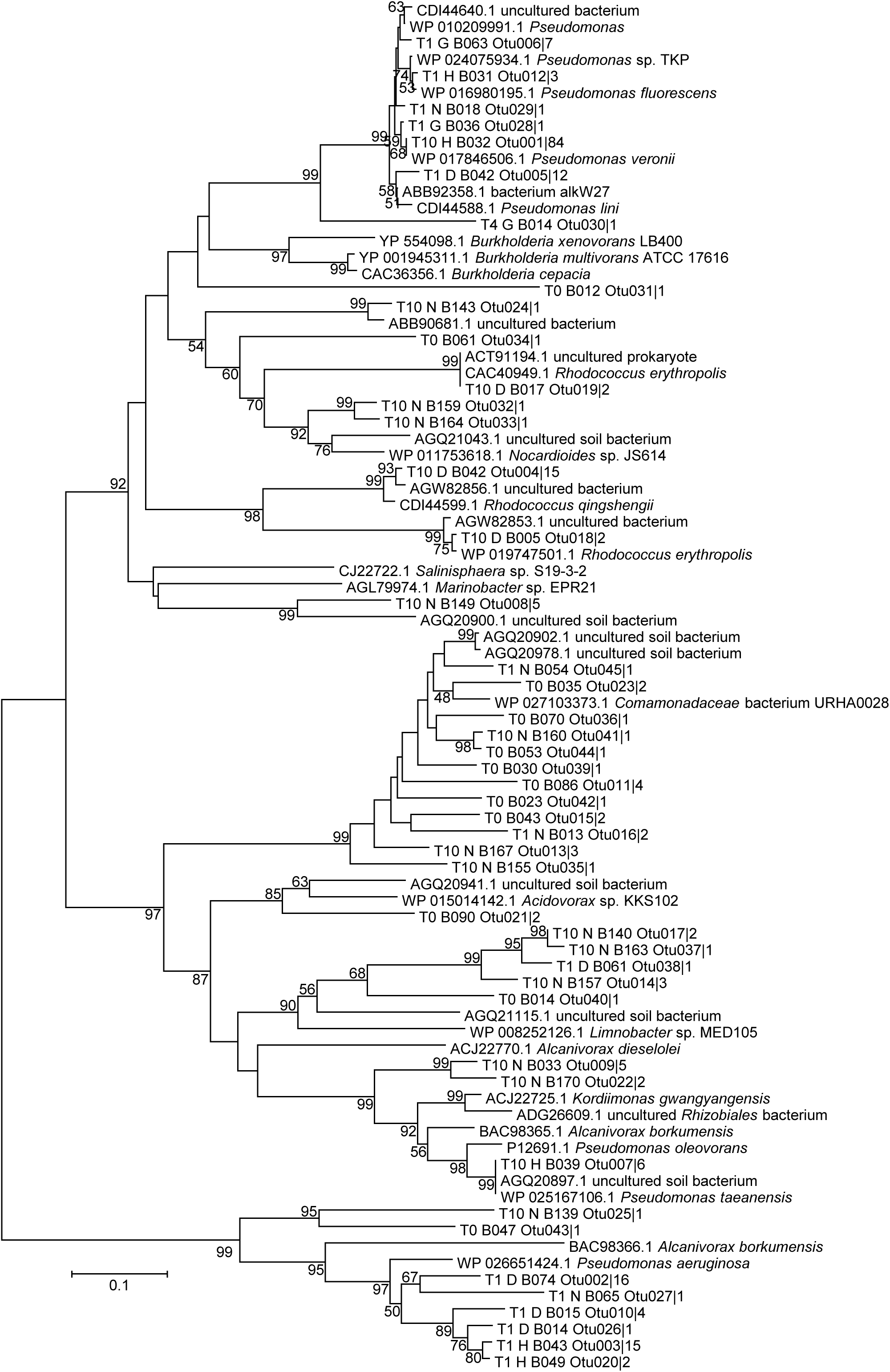
Phylogenetic distribution of *alkB* sequences. The phylogenetic tree was constructed from a ClustalW alignment of the derived amino acid sequences by neighbor-joining analysis with 1000 bootstrap replicates using Mega5.0 (Tamura et al. 2011). Bootstrap values greater than 50 are indicated.

Rarefaction curves (Supplemental Fig. 1A) constructed for each *alkB* library indicated that all libraries except the T0 and the 1-and 10-day unamended control libraries were adequately sampled (ie. plateaus were observed

#### *CYP 153A1* clone library analysis

*CYP 153A1* clone libraries were constructed from amplicons from the initial sediment sample (T0 NA), the no-alkane exposed control one and ten day samples (T1 NA and T10 NA), and the 1-and 10-day dodecane exposed samples (T1 D and T10 D), see Fig. 1B, 3 and Table 1. The other samples did not yield sufficient *CYP 153A1* PCR product for library construction. Furthermore, the T1 NA and the T1 and T10 D libraries contained very few sequences (Table 1). The T0 initial sediment sample contained sequences that assigned to 13 OTUs and had the highest Chao1 and Shannon Weaver scores (see Table 1). These sequences matched by BlastX to *CYP 153A1* from the alpha proteobacterial order *Rhizobiales*, and genera *Sphingopyxis*, and *Parvibaculum*, as well as several uncultured bacteria that group with the alphaproteobacteria (Fig. 3). The T1 NA library yielded only three sequences, each belonging to different OTUs; one *Rhizobiales-like* sequence (OTU6), a *Rhodococcus* (OTU27) and *Parvibaculum hydrocarboniclasticum* (OTU31). T10 NA contained predominantly *P. hydrocarboniclasticum* (OTU3) and *Rhizobiales* (OTU2). The only sequence recovered from the one-day dodecane (T1 D) exposed sample environment most closely matched a *CYP 153A1* from an uncultured *Rhizobiales* (OTU16) (Fig. 1B, Table 2). By 10 days of dodecane exposure most of the *CYP 153A1* sequences matched either *CYP 153A1* sequences from *Rhodococcus fasciens* (OTU1) or *Rhodococcus erythropolis* (OTU7).

**Figure 3.**
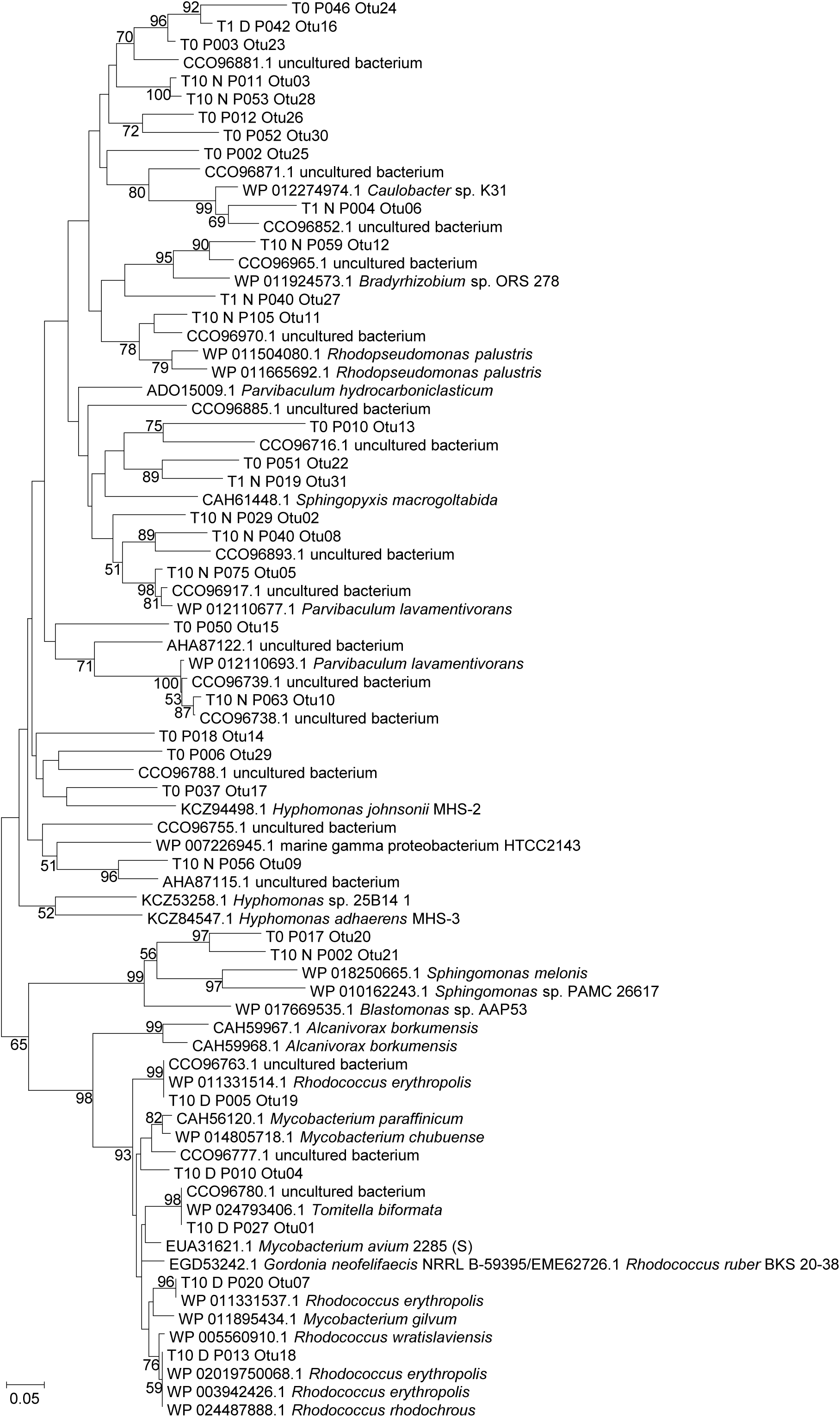
Phylogenetic distribution of *P450* sequences. The phylogenetic tree was constructed from a ClustalW alignment of the derived amino acid sequences by neighbor-joining analysis with 1000 bootstrap replicates using Mega5.0 (Tamura et al. 2011). Bootstrap values greater than 50 are indicated.

Analysis of rarefaction curves (Supplemental Fig. S1B) constructed for each *CYP 153A1* library indicated that only the 10-day exposure libraries (dodecane and no-alkane) reached a plateau and thus were adequately sampled. Overall, the limited number of sequences recovered limits our ability to analyze *CYP 153A1* alkane hydroxylases from the microbial community studied, although it is interesting that the T10 D sample yielded a *Rhodococcus-type CYP 153A1* alkane hydroxylase sequence, similar taxonomically to what was observed in the T10 D *alkB* clone library dataset.

### Quantification of relative *alkB* and *CYP153A1* gene abundance in dilution cultures

QPCR was used to assess the relative abundance of the dominant alkane hydroxylase OTUs from the sample libraries. Primers were designed to target alkB OTUs 1-5 and *CYP 153A1* OTUs 1,4 and 7 (Table 2) (primers are in Supplementary Table 1, results in Table 3). *AlkB* OTU5 was the only *alkB* OTU detected in the T0 sample DNA and was also detected in all samples except for the 10 day unamended control sample, suggesting that it may not be specifically induced by alkanes. The T1 H sample contained very elevated OTU2 levels, a sequence which matches an alkane hydroxylase from *Pseudomonas aeruginosa* (Table 2). OTUs 1 and 3 (also matches to *Pseudomonas-derived* alkane hydroxylases) were also detected in this sample, although at lower levels. Ten days of hexane exposure (T10 H) resulted in the decline of OTUs 2 and 3, and an approximately 7-fold increase in OTU 1. In the gasoline-exposed samples, OTUs 1, 3 and 5 were found at the same levels in the 1-day and 10-day samples. OTU2 was not detected in the 1-day sample but was present at relatively high abundance in the 10-day gasoline-exposed sample (T10 G). The 1-day dodecane exposed sample contained detectable *alkB* OTUs 1, 2, 3 and 5 and no detectable OTU4. By day 10, however, OTU4 dominated. Overall, the *alkB* qPCR analysis supported the clone library analysis, demonstrating a selection for *Pseudomonas*-type *alkBs* in response to *n-* hexane and gasoline treatment, and for *Rhodococcus*-type *alkB* s in response to dodecane treatment. The *CYP 153A1* qPCR analysis was largely unsuccessful, with no detection of any of the selected OTUs other than the OTUs 1, 4 and 7 in the 10-day dodecane-exposed sample which may suggest *CYP 153A1* type alkane hydroxylases were not important in alkane degradation in this system or because of the limited data, suggested that more extensive sampling and analysis may be required.

**Table 3.** QPCR analysis of selected alkane hydroxylase OTU DNA. Standard deviations are included in parentheses.

| OTU | Abundance of OTU as: |  |  | <u>Copy # of OTU DNA/ng gDNA</u> |  |  |  |  |  |
| --- | --- | --- | --- | --- | --- | --- | --- | --- | --- |
|  | Sample<sup>a</sup> |  |  | Copy # 16S rDNA/ng gDNA |  |  |  |  |  |
| *AlkB* |  |  |  |  |  |  |  |  |  |
|  | T0 | T1 NA | T10 NA | T1 D | T10 D | T1 H | T10 H | T1 G | T10 G |
| 1 | ND | ND | ND | 0.05<br>(±0.08) | 0.02<br>(±0.02) | 0.40<br>(±0.33) | 3.00<br>(±2.01) | 0.33<br>(±0.27) | 0.48<br>(±0.55) |
| 2 | ND | 1.30<br>(±0.83) | ND | 0.21<br>(±0.29) | ND | 1009<br>(±873.83) | 0.02<br>(±0.02) | ND | 71<br>(±113.85) |
| 3 | ND | 0.74<br>(±0.42) | ND | 2.36<br>(±1.18) | 0.64<br>(±0.14) | 2.88<br>(±1.49) | 0.59<br>(±0.18) | 0.04<br>(±0.01) | 0.08<br>(±0.00) |
| 4 | ND | ND | ND | ND | 0.57<br>(±1.79) | ND | ND | 5.95<br>(±14.61) | ND |
| 5 | 0.05<br>(±0.01) | 0.03<br>(±0.02) | ND | 0.02<br>(±0.02) | 0.03<br>(±0.01) | 0.06<br>(±0.04) | 0.01<br>(±0.01) | 0.03<br>(±0.01) | 0.02<br>(±0.03) |
| *CYP 153A1* |  |  |  |  |  |  |  |  |  |
| 1 | ND | ND | ND | ND | 0.47<br>(±0.68) | ND | ND | ND | ND |
| 4 | ND | ND | ND | ND | 0.10<br>(±0.23) | ND | ND | ND | ND |
| 7 | ND | ND | ND | ND | 0.31<br>(±0.81) | ND | ND | ND | ND |
| 16 | ND | ND | ND | ND | ND | ND | ND | ND | ND |

<sup>a</sup> Sample names are given at times of sampling followed by type of alkane where ND = not detected. T = time of exposure (days), NA = no alkanes (control), D = dodecane, H = *n*-hexane, G = gasoline.

### 16S rRNA sequence analysis

Changes in the bacterial community in response to different alkanes were examined by tag sequencing of the V6 region of the 16S rRNA using an Ion Torrent PGM. A total of 321,310 reads were obtained, of which 281,782 passed quality filtering and were mapped to individual barcodes using the QIIME pipeline. The number of sequences mapped to each treatment ranged from a high of 68,610 in the initial sediment (T0) sample to a low of 92 sequences in the day 10 unamended control treatment (T10 NA) (Table 4). The processed data were then used to calculate alpha (Shannon, Chao1, phylogenetic distances and PCoA analyses; Table 4 and Fig. 6) and beta (Unifrac analysis; Fig. 5) diversity metrics at a sampling depth of 5000 with T10 NA excluded due to the low read number (Table 4).

**Table 4.** Summary of IonTorrent PGM 16S rDNA V6 tag sequencing efforts and alpha diversity metrics. Standard deviations are included in parentheses.

| Library | Barcode | Processed sequences | OTUs observed | Chao1 | Shannon Index | Phylogenetic Distance |
| --- | --- | --- | --- | --- | --- | --- |
| T0 | ATCAG | 68,610 | 2132 | 4942 (216) | 10.0 (0.01) | 75 |
| T1 NA | GTGAG | 23,508 | 528 | 1551 (201) | 5.0 (0.1) | 34 |
| T1 H | GATCT | 45,242 | 426 | 1274 (145) | 4.4 (0.0) | 28 |
| T1 D | CGTCT | 13,730 | 1736 | 5274 (344) | 8.4 (0.0) | 61 |
| T1 G | CGACG | 62,669 | 346 | 921 (70) | 3.9 (0.0) | 23 |
| T10 NA | AGCAT | 92 | nd | nd | nd | nd |
| T10 H | TGTCA | 9,794 | 407 | 407 (38) | 2.5 (0.0) | 16 |
| T10 D | GCGAT | 32,439 | 782 | 2075 (246) | 5.5 (0.0) | 42 |
| T10 G | TCTGT | 25,689 | 872 | 872 (106) | 4.5 (0.0) | 22 |

Between 93.6 % and 99.5 % of the processed sequences could be classified below the bacterial root using the QIIME pipeline with the majority of sequences classified to the family or genus level. OTUs were divided into 13 groups for ease of analysis (Fig. 4), with each Group assigned to a genus, family or Class, as could be achieved by our analysis. Group I comprised the sequences that could not be classified and accounted for between 0.5 and 6.4 % of all sequences obtained. Group XIII represents all rare OTUs (ie. each sequence type is ≤ 1 % of the sequences from a given sample) pooled into Group XIII. The taxonomic annotation for the remaining groups is included in the Fig. 4 legend.

**Figure 4.**
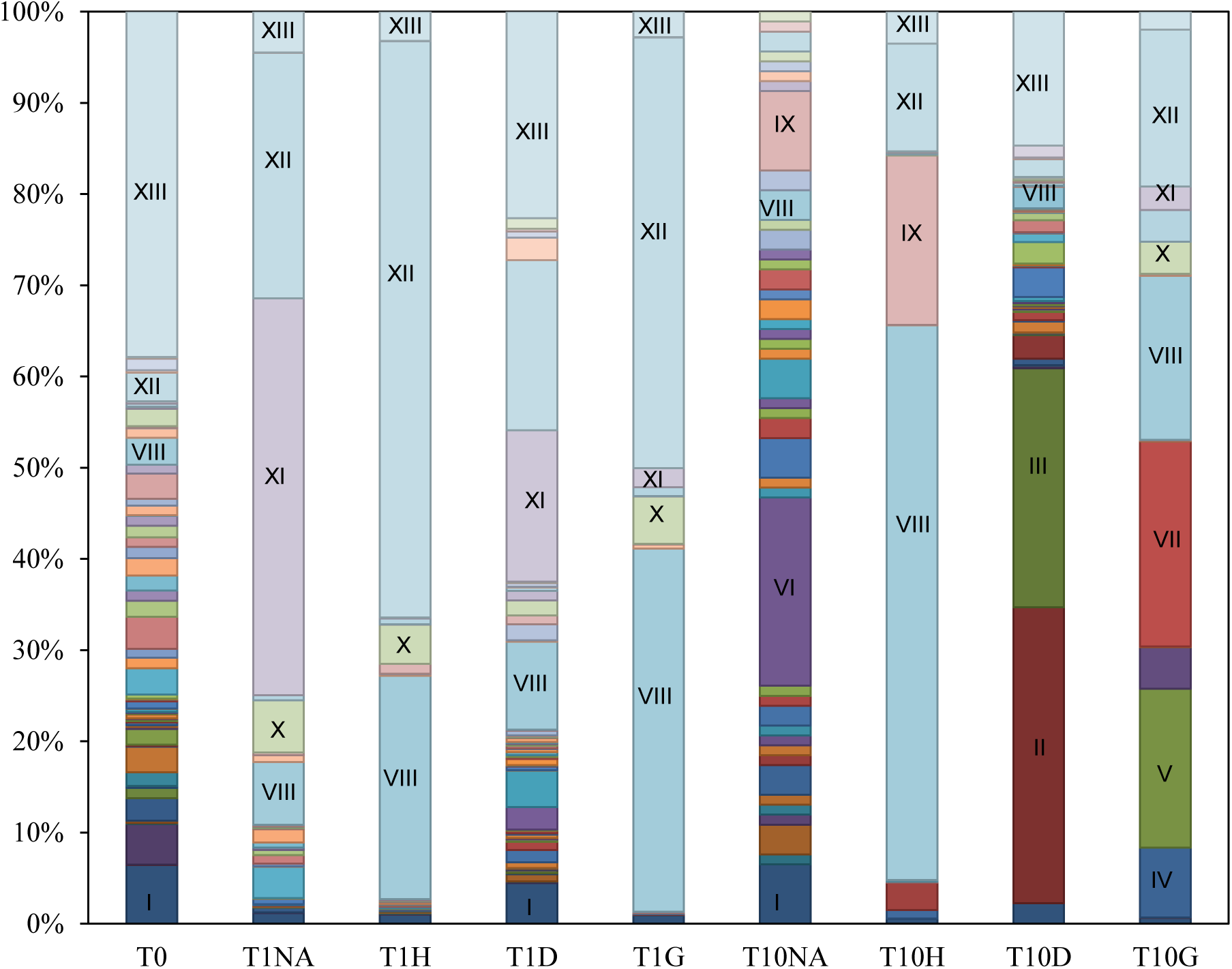
Bar graph showing distribution of tags into OTUs at the genus level. OTUs accounting for greater than 5% of an enrichment are labeled as follows: I, Other; II, Genus *Mycobacterium*; III, Genus *Rhodococcus*; IV, Genus *Bacillus*; V, Genus *Lysinibacillus*; VI, Family Aurantimonadaceae; VII, Family Acetobacteraceae; VIII, Class Gammaproteobacteria; IX, Genus *Marinobacter*; X, Genus *Shewanella*; XI, Genus *Acinetobacter*; XII, Genus *Pseudomonas*; XIII, groups that account for <1% of any enrichment.

The initial sediment bacterial community (T0) was the most diverse sample by all measures other than the Chao1 score (Table 4) with the most abundant OTU identified as Group XII *(Pseudomonas:* 3.2%). The T0 sample contained some sequences not found in any other sample, such as *Geobacter* (2.8 %), *Clostridium* (2.8 %), *Flavobacterium* (2.4 %) and *Burkholderiales* (10%). The Gram positive sequences were diverse and represented 6.5% of the total. Dilution alone (culture in the absence of alkane addition), reduced diversity and altered the community composition as seen in other sections (see Discussion). The T1 NA sample was less diverse than the T0 sample by all measures calculated (Table 4) and contained a community dominated by Gammaproteobacteria (84.7%) mainly *Acinetobacter* (Group XI: 43.5 %) and *Pseudomonas* (Group XII: 26.9%). The T10 NA sample was dominated by Gammaproteobacteria (22.8%), of which 8.7% were Group IX*Marinobacter*. This sample could not be assessed for diversity metrics as it contained only 92 reads.

While our assessment of community diversity is limited by the lack of ability to calculate diversity measures on the 10-day no alkane control (T10 NA), the one day analysis indicates that *n-* hexane and gasoline reduced the diversity of the communities sampled beyond the loss of diversity seen in one day of dilution alone (the no alkane control, T1 NA). Interestingly, this was not the case with dodecane exposure, which instead resulted in an increase in diversity by all measures after one day of exposure (T1 D). Comparing the diversity measures in all three types of alkane exposure after 10 days of exposure (T10 D, T10 H and T10 G) indicates that diversity continued to decline in the *n-* hexane and gasoline exposed treatments while it declined to a lesser degree in dodecane exposure. Hierarchical clustering (Fig. 5) from unweighted and weighted unifrac analyses revealed the dodecane-exposed samples clustered with the initial T0 sample, separate from all other samples. PCoA analysis (Fig. 6) revealed that the similarities between the different hydrocarbon amendments and the no-hydrocarbon control dilution culture were driven mainly by the *Gammaproteobacteria*.

One day of exposure to *n-* hexane and gasoline shifted the community to predominantly Gammaproteobacteria, especially *Pseudomonas* (Group XII) and Gammaproteobacteria (Group VIII). This Gammaproteobacterial dominance persisted to 10 days in the *n-* hexane exposed culture, but in the gasoline-exposed culture the proportion of *Gammaproteobacteria* declined by 10 days and an increase in *Alphaproteobacteria*, specifically the family *Acetobacteraceae* (Group VII: 22.5%) occurred. The proportion of Gram positive bacteria increased in the T10 G sample to 29.7%, predominantly Firmicutes from the genera *Bacillus, Lysinibacillus* and *Rumelibacillus*. As observed in the analysis of *alkB* clone sequences, dodecane exposure caused a transient shift towards Gammaproteobacteria (56.2%) after one day of exposure but by 10 days of continued dodecane exposure the *Gammaproteobacteria* declined (to 3.1% of sequences), and an upsurge in Gram positive bacteria (to 59.0%) occurred. The Gram positive bacteria were mainly from the class *Actinobacteria*, chiefly *Mycobacterium* (Group II: 32.4 %) and *Rhodococcus* (Group III: 26.2 %). Overall, the T10 D sample contained the greatest proportion of Gram positive sequences observed in any sample (The T0 NA sample contained 6.5%, all T1 samples contained less than one percent, T10 H contained 3.9%, T10 G 29.7% and T10 D 59.0%).

**Figure 5.**
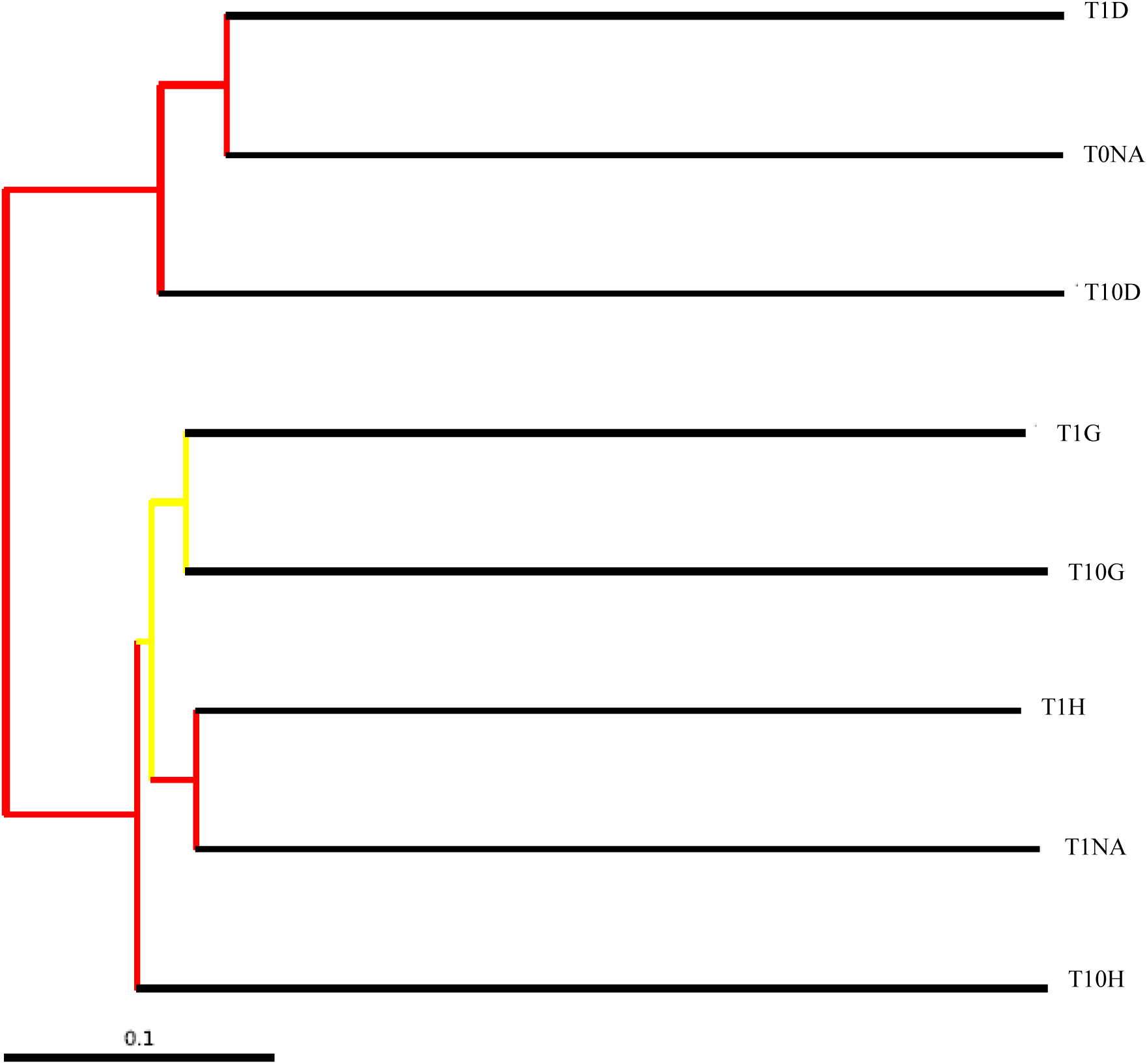
Dendrograms of hierarchical clustering of community similarities from all libraries (excluding the T10 NA due to low number of sequences recovered) based on weighted UniFrac Jackknife Cluster Analysis with 100 permutations (5000 sequence sub-sampling). Each node is colored by the fraction of times it was recovered in the jackknife replicates. Nodes recovered >99.9% of the time are red, 90-99.9% are yellow. T0, T1 and T10 indicate zero (initial), 1 day and 10 day enrichment samples. Alkanes are indicated as D (dodecane), H (hexane), G (gasoline) or NA (no alkane control).

**Figure 6.**
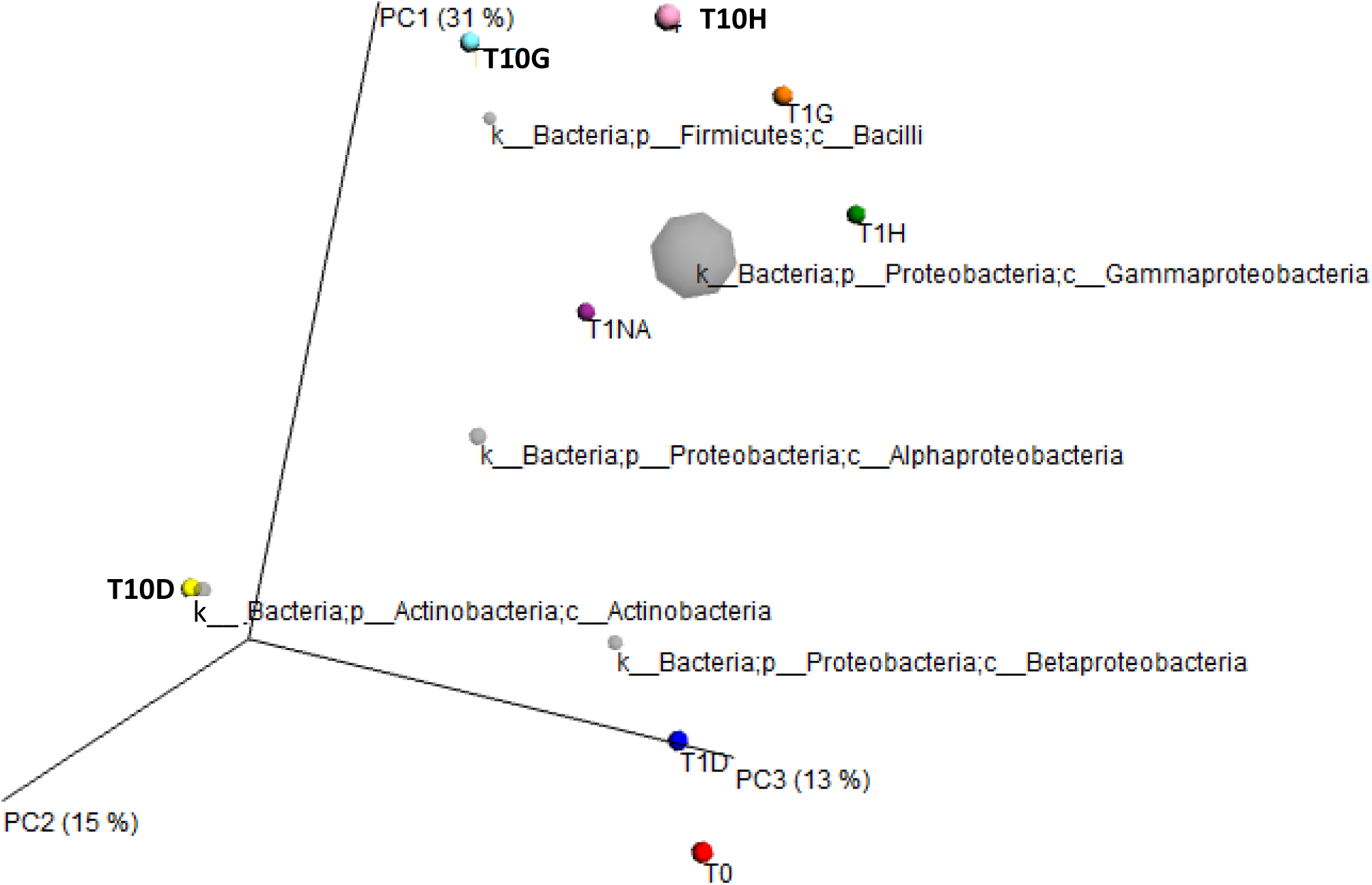
Principal coordinate analysis (PCoA) ordination based on an unweighted UniFrac distance matrix of sediment bacterial 16S rDNA gene profiles sampled from the initial sediment (T0), and after 1 and 10 days of exposure to dodecane (D), hexane (H) and gasoline (G) and no alkane control (NA). The T10 day no alkane control sample was not included due to the low number of sequences recovered. K ordination value labels indicate p = phylum and c = class.

In total, the analysis of the microbial community agrees with the *alkB* clone library analysis, indicating that Gammaproteobacteria are important members of the microbial community that respond quickly to both dilution and alkane exposure, although the specific chemical nature of the alkane present influences the community response. The ability of gasoline, and dodecane exposure, to recruit Gram positive response is also noted.

## Discussion

Small brackish estuarine tidal marshes, referred to as fringing marshes, are an important coastal habitat in New Hampshire (Morgan et al., 2009 and references therein, PREP 2013), providing essential ecosystem services similar to larger meadow marshes. These fringing marshes are under pressure from sea level rise and exposure to pollutants such as petroleum (Morgan et al., 2009; NHDES 2004; PREP 2013; Stralberg et al., 2011). Unlike larger meadow marshes, relatively little is known about the ability of fringing marsh microbial communities to degrade petroleum hydrocarbons, knowledge that would help us better predict marsh resilience. In this study. we assessed the *n-* alkane degrading potential of a fringing marsh microbial community from the Cocheco River of the Great Bay of New Hampshire, a site where the role of *Spartina alterniflora* in phytoremediation of PAHs has been documented (Watts et al., 2006).

By focusing on alkane hydroxylase genes *(alkB* and *CYP 153A1)* the functional diversity was assessed in the initial sediment, and dilution cultures treated with *n-* hexane, gasoline, dodecane or dilution alone (no alkanes) for 10 days. The initial community was already enriched in *Pseudomonas*-type *alkB* sequences which increased in relative abundance by one day’s exposure to dilution alone, and all three sources of alkanes tested. The initial dominance and rapid expansion of Gammaproteobacteria, especially *Pseudomonas spp.* has been shown in a variety of environments including meadow salt marshes (Acosta-Gonzales et al., 2015; Atlas et al., 2015; Beazley et al., 2012; Kasai et al., 2001; Koo et al., 2015; Kostka et al., 2011; Lamendella et al., 2014; Liu & Liu 2013, Looper et al., 2013). By 10 days of further culture we observed a differential response according to treatment condition, with gasoline and *n-* hexane further enriching the fraction of *Pseudomonas*-type *alkBs* and reducing the overall diversity of *alkBs* recovered to a greater extent than that seen in the no alkane control or the dodecane exposed cultures. In contrast, dodecane exposure enriched for *Rhodococcus-like alkB* sequences and resulted in a 10-day community with slightly more diverse *alkBs* than seen in the no alkane control. Interestingly we were only able to amplify *CYP 153A1*-type genes from the no-alkane control and dodecane dilution cultures and failed to detect *CYP 153A1* genes in most samples using QPRC (Table 1, Supplemental Table 1). This may suggest that *CYP 153A1* alkane hydroxylases are not as important as *alkB* type hydroxylases in the fringing marsh we studied, which contrasts with other reports focusing on petroleum-degrading bacteria in marine and soil environments (van Beilen et al., 2006, Wang et al., 2010a). The role (or lack of role) of CYP 153A1 type alkane hydroxylases in fringing salt marsh microbial communities requires further study.

Analysis of the 16S rRNA V6 region to assess the phylogenetic diversity and taxonomic composition of the whole community generally supported the findings of the clone library functional analysis. The initial sediment community was highly diverse, with the dominant taxa present (3.2%) matching to *Pseudomonas* spp. All dilution cultures, including the no-alkane control, exhibited a decrease in diversity, richness and phylogenetic distance (PD) after one day of dilution culture (Fig. 4, Table 4) which is expected given that dilution alone reduces diversity (Franklin et al., 2001, Hewson et al., 2003, Schaffer et al., 2000, Yan et al., 2015). Comparing the diversity values obtained after one day of treatment (we were not able to compare diversity values at 10 days of treatment due to the low read number in the 10-day no alkane control), and the 10 day cultures, indicated that exposure to gasoline and *n-* hexane reduced diversity to a greater degree than seen in the dodecane treatment or the no alkane control treatment. This agrees with our functional gene analysis and the findings of other studies documenting the negative impact of highly concentrated petroleum hydrocarbons on microbial diversity (Acosta-González et al., 2015; Atlas et al., 2015; Beazley et al., 2012; Guibert et al., 2012; Mason et al., 2012; Wang et al., 2011; Pérez-de-Mora et al., 2011; Saul et al., 2005). Others have reported a rebound in diversity once petroleum hydrocarbon concentrations decline (through degradation: Atlas et al., 2015; Koo et al., 2015; Paisse et al., 2010). Given that we maintained saturating concentrations of n-alkanes, further study would be required to determine if the sediment microbial community of the fringing marsh we studied is as resilient as larger marshes appear to be in the face of petroleum influx (Acosta-González et al., 2015; Atlas et al., 2015 and others).

The taxonomic composition of the entire microbial community varied by treatment, generally supporting the taxonomic assignment of *alkB* genes recovered. Gammaproteobacteria, particularly the rapidly responding *Pseudomonads*, dominated in the initial sediment and were enriched in all one day dilution cultures (alkane-amended or unamended) (Fig. 4), as well as in the 10 day gasoline and *n*-hexane exposed cultures. Gammaproteobacteria are known to respond rapidly to dilution alone (Yan et al., 2015), as well as being important early responders in petroleum-contaminated sites including larger salt marshes (references above).

Interestingly, by 10 days of exposure to dodecane and gasoline, there was an increased proportion of Gram positive bacteria, with the T10 D sample containing 59.0% Gram positive sequences mainly from the genera *Rhodococcus* (26.2%) and *Mycobacterium* (32.4%), while the T10 G sample was dominated by Firmicutes (29.7%), mainly from the genera *Bacillus, Lysinibacillus* and *Rumelibacillus*. *Rhodococcus* species are known alkane degraders (Hassanshahian et al., 2010, Larkin et al., 2005, Liu & Liu, 2013, Sekine et al., 2006, Van Hamme & Ward 2001, Whyte et al., 1998 & 2002, Xu et al., 2007) documented in petroleum-impacted salt marshes (Acosta-González et al., 2015; Mahmoudi et al., 2013, Liu & Liu 2013), as are Mycobacterium (Atlas et al., 2015 and others). The role of Gram positive bacteria in petroleum degradation has been established in larger salt marshes and other systems (Acosta-González et al., 2015; Atlas et al., 2015; Koo et al., 2015; Launen et al., 2008; Looper et al., 2012; Liu & Liu 2013; Mahmoudi et al., 2013) and there is evidence that Gram positive bacteria may be important in the secondary response to petroleum degradation, increasing in abundance more slowly than the rapidly responding Gammaproteobacteria (Koo et al., 2015). Our results suggest that the role of Gram positive bacteria in responding to alkanes in fringing marshes may be similar.

Selection of certain microbial community members over others when challenged with different types of n-alkanes may reflect differences in alkane-metabolizing enzymes, or physiological differences that affect the update or other aspects of alkane metabolism between microbes (Grund et al., 1975; Larkin et al., 2005; Smits et al., 2002; Van Beilen et al., 1994; Van Beilen et al., 2005; Wang & Shao 2013; Wentzel et al., 2007), alkane toxicity (Ramos et al., 2002; Vermuë et al., 1993; Walker & Colwell 1975), alkane uptake from the environment (Bouchez-Naïtali et al., 1999; Rosenberg & Rosenberg 1981; Van Hamme & Ward 2001), biosurfactants (Desai & Banat 1997; Ivshina et al., 1998; Pornsunthorntawee et al.,2008, Pi et al.,2017) or relative growth rates (Vieira-Silva & Rocha 2010). Resolving the mechanisms underlying the selection for specific community members by specific alkanes, as we observed here, requires further study, likely involving a culture-based approach.

Overall, our results indicate that the fringing marsh studied contains a diverse microbial community enriched with *Pseudomonas-type* alkane degrading bacteria able to expand rapidly when exposed to environmental change (dilution and alkane exposure), as has been observed in larger meadow-type salt marshes elsewhere (Acosta-González et al., 2015; Beazley et al., 2012; Hazen et al., 2010; Mason et al., 2012). The importance of Gram positive bacteria, especially *Rhodococcus*, in a slower response to dodecane exposure was also observed, and agrees with findings in meadow marshes and other environments (Acosta-Gonzâlez et al., 2015; Atlas et al., 2015; Mahmoudi et al 2013, Liu & Liu 2013). Overall, our data suggest that the microbial community studied is capable of responding to perturbation by alkanes, and that it does so in an alkane-specific manner, supporting the potential of such fringing marshes to buffer against petroleum influx through degradation. Given the importance of fringing marshes in the New England region, and the threat of petroleum influx into these marshes, it seems likely that more study is merited. Two areas of particular relevance would be an evaluation of the ability of these communities to degrade whole-oil mixtures, such as home heating oil (which is regularly transported in the Great Bay Estuary), as well as an assessment of the ability of these communities to recover diversity once petroleum has been degraded.

## Acknowledgements

We thank Ms. Katie Featherston, Ms. Marianne O’Brien, Ms. Audrey Arsenault, Dr. Penny Micele and Dr. Gordon Leversee for support in the administration of this work.

## References

Acosta-González, A., Martirani-Von Abercron, S. M., Rosselló-Móra, R., Wittich, R. M., and Marqués, S. (2015). The effect of oil spills on the bacterial diversity and catabolic function in coastal sediments: a case study on the Prestige oil spill. Environ. Sci. Pollut. Res. Int. 22, 15200–15214. doi:10.1007/s11356-015-4458-y

Asperger O, Naumann A & Kleber HP (1981) Occurrence of cytochrome P450 in Acinetobacter strains after growth on n-hexadecane. FEMS Microbiol Lett 11: 309–12. doi:10.1111/j.W1574-6968.1981.tb06986.

Atlas RM (1975) The effects of temperature and crude oil composition on petroleum bioremediation. Appl. Microbiol. 33 (3):396–403

Atlas RM & Hazen TC (2011) Oil biodegradation and bioremediation: a tale of the two worst spills in U.S. history. Environ Sci Technol 45: 6709–15. doi:10.1021/es2013227

Atlas RM, Stoeckel DM, Faith SA, Minard-Smith A, Thorn JR & Benotti MJ (2015) Oil biodegradation and oil-degrading microbial populations in marsh sediments impacted by oil from t he deepwater horizon well blowout. Environ. Sci. Technol. 49 (14):8356–8366. doi:10.1021/acs.est.5b00413

Beazley MJ, Martinez RJ, Rajan S et al.,(2012) Microbial community analysis of a coastal salt marsh affected by the Deepwater Horizon oil spill. PLoS One 7: e41305. doi:10.1371/journal.pone.0041305. Epub 2012 Jul 18.

Boopathy, R., Shields, S., and Nunna, S. (2012). Biodegradation of crude oil from the BP oil spill in the marsh sediments of southeast Louisiana, USA. Appl. Biochem. Biotechnol. 167, 1560–1568. doi:10.1007/s12010-012-9603-1

Bouchez-Naïtali M, Rakatozafy H, Marchal R, Leveau JY& Vandecasteele JP (1999) Diversity of bacterial strains degrading hexadecane in relation to the mode of substrate uptake. J Appl Microbiol 86: 421–8. doi:10.1046/j.1365-2672.1999.00678.x

Brown DG, Gupta L, Kim TH, Moo-Young HK & Coleman AJ (2006) Comparative assessment of coal tars obtained from 10 former manufactured gas plant sites in the eastern United States. Chemosphere 65: 1562–9. doi:10.1016/j.chemosphere.2006.03.068

Caporaso J, Kuczynski J, Stombaugh J et al.,(2010a) QIIME allows analysis of high-throughput community sequencing data. Nat Methods 7: 335–6. doi:10.1038/nmeth.f.303

Caporaso JG, Bittinger K, Bushman FD, DeSantis TZ, Andersen GL & Knight R (2010b) PyNAST: a flexible tool for aligning sequences to a template alignment. Bioinformatics 26: 266–7. doi:10.1093/bioinformatics/btp636. Epub 2009 Nov 13.

Chao A (1984) Nonparametric estimation of the number of classes in a population. Scand J Stat 11: 265–70.

Chao A & Lee S-M (1992) Estimating the number of classes via sample coverage. J Am Stat Assoc 87: 210–7.

Desai JD & Banat IM (1997) Microbial production of surfactants and their commercial potential. Microbiol Mol Biol R 61: 47–64.

Felsenstein J (2005) PHYLIP (Phylogeny Inference Package) version 3.6. Department of Genome Sciences, University of Washington, Seattle., Distributed by the author.

Franklin R, Garland J, Bolster C, Mills A. (2001) Impact of dilution on microbial community structure and functional potential: comparison of numerical simulation and batch culture experiments. Appl Environ Microbiol. 67: 702–712. doi:10.1128/AEM.67.2.702-712.2001

Grund A, Shapiro J, Fennewald M, Bacha P, Leahy J, Markbreiter K, Nieder M & Toepfer M (1975) Regulation of alkane oxidation in Pseudomonas putida. J Bacteriol 123: 546–56.

Guibert L, Loviso C, Marcos M, Commendatore M, Dionisi H & Lozada M (2012) Alkane biodegradation genes from chronically polluted subantarctic coastal sediments and their shifts in response to oil exposure. Microb Ecol 64: 605–16. doi:10.1007/s00248-012-0051-9

Hassanshahian M, Emtiazi G, Kermanshahi RK, et al., (2010) Comparison of oil degrading microbial communities in sediments from Persian Gulf and Caspian Sea. Soil Sediment Contam. 19: 277–291. doi:10.1080/15320381003695215

Hewson I., Vargo GA & JA Fuhrman (2003). Bacterial diversity in shallow oligotrophic marine benthos and overlying waters: effects of virus infection, containment, and nutrient enrichment. Microb. Ecol. 46, 322–336. doi:10.1007/s00248-002-1067-3

Huber JA,Welch DBM, Morrison HG, Huse SM, Neal PR, Butterfield DA & Sogin ML (2007) Microbial population structures in the deep marine biosphere. Science 318: 97–100. doi:10.1126/science.1146689

Ivshina IB, Kuyukina MS, Philp JC & Christofi N (1998) Oil desorption from mineral and organic materials using biosurfactant complexes produced by Rhodococcus species. World J Microb Biot 14: 711–7. doi:10.1023/A:1008885309221

Kasai Y, Kishira H, Syutsubo K, Harayama S (2001) Molecular detection of marine bacterial populations on beaches contaminated by the Nakhodka tanker oil-spill accident. Environ Microbiol 3:246–55. doi:10.1046/j.1462-2920.2001.00185.x

Kimes NE, Callaghan AV, Suflita JM & Morris PJ (2014) Microbial transformation of the Deepwater Horizon oil spill –past, present, and future perspectives. Front Microbiol 18; 5:603. doi:10.3389/fmicb.2014.00603. eCollection 2014.

Kloos K, Munch JC & Schloter M (2006) A new method for the detection of alkane-monooxygenase homologous genes (alkB) in soils based on PCR-hybridization. J Microbiol Meth 66: 486–96. doi:10.1016/j.mimet.2006.01.014

Kok M, Oldenhuis R, van der Linden MP, Raatjes P, Kingma J, van Lelyveld PH & Witholt B (1989) The Pseudomonas oleovorans alkane hydroxylase gene. Sequence and expression. J Biol Chem 264: 5435–41.

Koo, H.; Mojib, N.; Thacker, R.W.; Bej, A.K. (2014) Comparative analysis of bacterial community metagenomics in coastal Gulf of Mexico sediment microcosms following exposure to Macondo oil (MC252). Antonie Van Leeuwenhoek 106 (5):993–1009. doi:10.1007/s10482-014-0268-3

Kostka JE, Prakash O, Overholt WA, Green SJ, Freyer G, Canion A, Delgardio J, Norton N, Hazen TC, Huettel M (2011) Hydrocarbon-degrading bacteria and the bacterial community response in Gulf of Mexico beach sands impacted by the Deepwater Horizon oil spill. Appl Environ Microbiol 77:7962–7974. doi:10.1128/AEM.05402-11

Kubota M, Nodate M, Yasumoto-Hirose M, Uchiyama T, Kagami O, Shizuri Y & Misawa N (2005) Isolation and functional analysis of cytochrome P450 CYP153A genes from various environments. Biosci Biotechnol Biochem 69: 2421–30. doi:10.1271/bbb.69.2421

Lamendella R, Strutt S, Borglin S, Chakraborty R, Tas N, Mason OU, Hultman J, Prestat E, Hazen TC, Jansson JK (2014) Assessment of the Deepwater Horizon oil spill impact on gulf coast microbial communities. Front Microbiol 5:130. doi:10.3389/fmicb.2014.00130

Larkin MJ, Kulakov LA & Allen CCR (2005) Biodegradation and Rhodococcus –masters of catabolic versatility. Curr Opin Biotech 16: 282–90. doi:10.1016/j.copbio.2005.04.007

Launen LA, Dutta J, Turpeinen R, Eastep ME, Dorn R, Buggs VH, Leonard JW & Häggblom MM (2008) Characterization of the indigenous PAH-degrading bacteria of Spartina dominated salt marshes in the New York/New Jersey Harbor. Biodegradation 19: 347–63. doi:10.1007/s10532-007-9141-7

Liu Z, & J Liu (2013). Evaluating bacterial community structures in oil collected from the sea surface and sediment in the northern Gulf of Mexico after the Deepwater Horizon oil spill. MicrobiologyOpen 2, 492–504. doi:10.1002/mbo3.89

Liu C, Wang W, Wu Y, Zhou Z, Lai Q & Shao Z (2011) Multiple alkane hydroxylase systems in a marine alkane degrader, Alcanivorax dieselolei B-5. Environ Microbiol 13: 1168–78. doi:10.1111/j.1462-2920.2010.02416.x

Looper JK, Cotto A, Kim B-Y, Lee M-K, Liles MR, NíChadhain SM & Son A (2013) Microbial community analysis of Deepwater Horizon oil-spill impacted sites along the Gulf coast using functional and phylogenetic markers. Env Sci Process Impact 15: 2068–79. doi:10.1039/c3em00200d

Lozupone C & Knight R (2005) UniFrac: a new phylogenetic method for comparing microbial communities. Appl Environ Microbiol 71: 8228–35. doi:10.1128/AEM.71.12.8228-8235.2005

Lu Z., Deng Y., Van Nostrand J. D., He Z., Voordeckers J., Zhou A., et al. (2012). Microbial gene functions enriched in the Deepwater Horizon deep-sea oil plume. ISME J. 6: 451–460. doi:10.1038/ismej.2011.91

Magnusson M, C Colgan & Gittell R. (2012). The economic impact of the Piscataqua River and the Ports of Portsmouth and Newington. State of New Hampshire Division of Ports and Harbors. http://www.portofnh.org/documents/port_study_mm_6_7_12FINAL.pdf

Mahmoudi, N.; Porter, T.M.; Zimmerman, A.R.; Fulthorpe, R.R.; Kasozi, G.N.; Silliman, B.R.; Slater, G.F. (2013) Rapid degradation of Deepwater Horizon spilled oil by indigenous microbial communities in Louisiana saltmarsh sediments. Environ. Sci. Technol. 47 (23):13303–13312. doi:10.1021/es4036072

Maier T, Forster HH, Asperger O & Hahn U (2001) Molecular characterization of the 56-kDa CYP153 from Acinetobacter sp. EB104. Biochem Biophys Res Commun 286: 652–8. doi:10.1006/bbrc.2001.5449

Makepeace DK, Smith DW & Stanley SJ (1995) Urban stormwater quality: Summary of contaminant data. Crit Rev Env Sci Tech 25: 93–139. doi:10.1080/10643389509388476

Mason OU, Hazen TC, Borglin S, Chain PS, Dubinsky EA, Fortney JL, Han J, Holman HY, Hultman J, Lamendella R, Mackelprang R, Malfatti S, Tom LM, Tringe SG, Woyke T, Zhou J, Rubin EM, Jansson JK (2012) Metagenome, metatranscriptome and singlecell sequencing reveal microbial response to Deepwater Horizon oil spill. ISME J 6:1715–27. doi:10.1038/ismej.2012.59.

McGenity T. J. (2014). Hydrocarbon biodegradation in intertidal wetland sediments. Curr. Opin. Biotechnol. 27: 46–54. 10.1016/j.copbio.2013.10.010

Morgan, Pamela A.; Burdick, David M.; and Short, Frederick T., The Functions and Values Of Fringing Salt Marshes In Northern New England, USA (2009). Estuaries and Coasts 32(3):483–495. doi:10.1007/s12237-009-9145-0

Mukherjee S, Juottonen H, Siivonen P, Lloret C, Quesada CL, Tuomi P, Pulkkinen P & Yrjälä K. (2014) Spatial patterns of microbial diversity and activity in an aged creosote-contaminated site. ISME J. 8(10): 2131–2142. doi:10.1038/ismej.2014.151

Natter M, Keevan J, Wang Y, Keimowitz AR, Okeke BC, Son A, Lee MK. (2012) Level and degradation of Deepwater Horizon spilled oil in coastal marsh sediments and pore-water. Environ. Sci. Technol. 46 (11): 5744–5755. doi:10.1021/es300058w

NHDES, 2004. Environmental Fact Sheet WMB-CP-08. Threats to The Salt Marsh Environment. Available at: https: http://www.des.nh.gov/organization/commissioner/pip/factsheets/cp/documents/cp08.pdf

NHDES. 2015. State of New Hampshire DRAFT 2014 Section 303(d) Surface Water Quality List. NHDES-R-WD-15-11. October 14, 2015. New Hampshire Department of Environmental Services, Concord, NH.

Paisse S, Goni-Urriza M, Coulon F & Duran R (2010) How a bacterial community originating from a contaminated coastal sediment responds to an oil input. Microb Ecol 60: 394–405. doi:10.1007/s00248-010-9721-7

Pérez-de-Mora A, Engel M & Schloter M (2011) Abundance and diversity of n-alkane-degrading bacteria in a forest soil co-contaminated with hydrocarbons and metals: a molecular study on alkB homologous genes. Microbial Ecol 62: 959–72. doi:10.1007/s00248-011-9858-z

Pielou EC (1977) Mathematical ecology. New York: Wiley.

Pornsunthorntawee O, Wongpanit P, Chavadej S, Abe M & Rujiravanit R (2008) Structural and physicochemical characterization of crude biosurfactant produced by Pseudomonas aeruginosa SP4 isolated from petroleum-contaminated soil. Bioresource Technol 99: 1589–95. doi:10.1016/j.biortech.2007.04.020

Piscataqua River Estuaries Partnership (PREP) State of Our Estuaries (2013) University of New Hampshire, Dover, NH. pp. 48

Ramos JL, Duque E, Gallegos MT, Godoy P, Ramos-González MI, Rojas A, Teran W & Segura A (2002) Mechanisms of solvent tolerance in gram-negative bacteria. Annu Rev Microbiol 56: 743–68. doi:10.1146/annurev.micro.56.012302.161038

Rojo F (2009) Degradation of alkanes by bacteria. Environ Microbiol 11: 2477–90. doi:10.1111/j.1462-2920.2009.01948.x

Rosenberg M & Rosenberg E (1981) Role of adherence in growth of Acinetobacter calcoaceticus RAG-1 on hexadecane. J Bacteriol 148: 51–7.

Saul DJ, Aislabie JM, Brown CE, Harris L & Foght JM (2005) Hydrocarbon contamination changes the bacterial diversity of soil from around Scott Base, Antarctica. FEMS Microbiol Ecol 53: 141–55. doi:10.1016/j.femsec.2004.11.007:

Schafer, H., Servais, P., and Muyzer, G. (2000). Successional changes in the genetic diversity of a marine bacterial assemblage during confinement. Arch. Microbiol. 173, 138–145. doi:10.1007/s002039900121

Schloss P & Handelsman J (2009) Open-source, platform-independent, community-supported software for describing and comparing microbial communities. Appl Environ Microbiol 75: 7537–41. doi:10.1128/AEM.01541-09

Schneiker S, dos Santos VAPM, Bartels D et al., (2006) Genome sequence of the ubiquitous hydrocarbon-degrading marine bacterium Alcanivorax borkumensis. Nat Biotech 24: 997–1004. doi:10.1038/nbt1232

Sekine M, Tanikawa S, Omata S et al., (2006) Sequence analysis of three plasmids harboured in Rhodococcus erythropolis strain PR4. Environ Microbiol 8: 334–46. doi:10.1111/j.1462-2920.2005.00899.x

Smits THM, Balada SB, Witholt B & van Beilen JB (2002) Functional analysis of alkane hydroxylases from Gram-negative and Gram-positive bacteria. J Bacteriol 184: 1733–42. doi:10.1128/JB.184.6.1733-1742.2002

Speight JG (1998) The Chemistry and Technology of Petroleum: 3 Edition. New York, NY, USA: Marcel Dekker Inc.

Stralberg D, Brennan M, Callaway JC, Wood JK, Schile LM, et al., (2011) Evaluating tidal marsh sustainability in the face of sea–level rise: A hybrid modeling approach applied to San Francisco Bay. PLoS One 6 (11):e27388 doi:10.1371/journal.pone.0027388

Tamura K, Peterson D, Peterson N, Stecher G, Nei M & Kumar S (2011) MEGA5: molecular evolutionary genetics analysis using maximum likelihood, evolutionary distance, and maximum parsimony methods. Mol Biol Evol 10: 2731–9. doi:10.1093/molbev/msr121

van Beilen JB & Funhoff EG (2007) Alkane hydroxylases involved in microbial alkane degradation. Applied Microbiol Biotechnol 74: 13–21. doi:10.1007/s00253-006-0748-0

van Beilen JB, Funhoff EG, van Loon A et al., (2006) Cytochrome P450 alkane hydroxylases of the CYP153 family are common in alkane-degrading eubacteria lacking integral membrane alkane hydroxylases. Appl Environ Microbiol 72: 59–65. doi:10.1128/AEM.72.1.59-65.2006

van Beilen JB, Kingma J & Witholt B (1994) Substrate specificity of the alkane hydroxylase system of Pseudomonas oleovorans GPo1. Enzyme Microb Tech 16: 904–11. doi:10.1016/0141-0229(94)90066-3

van Beilen JB, Smits THM, Roos FF, Brunner T, Balada SB, Röthlisberger M & Witholt B (2005) Identification of an amino acid position that determines the substrate range of integral membrane alkane hydroxylases. J Bacteriol 187: 85–91. doi:10.1128/JB.187.1.85-91.2005

van Beilen JB, Smits THM, Whyte LG, Schorcht S, Röthlisberger M, Plaggemeier T, Engesser K-H & Witholt B (2002) Alkane hydroxylase homologues in Gram-positive strains. Environ Microbiol 4: 676–82. doi:10.1046/j.1462-2920.2002.00355.x

van Beilen JB, Veenhoff L & Witholt B (1998) Alkane hydroxylase systems in Pseudomonas aeruginosa strains able to grow on n-octane. In: Kieslich K, van der Beek CP, de Bont JAM, van den Tweel WJJ (eds.) Studies in Organic Chemistry volume 53: Elsevier, 211–5.

van Hamme JD & Ward OP (2001) Physical and metabolic interactions of Pseudomonas sp. strain JA5-B45 and Rhodococcus sp. strain F9-D79 during growth on crude oil and effect of a chemical surfactant on them. Appl Environ Microbiol 67: 4874–9. doi:10.1128/AEM.67.10.4874-4879.2001

Vazquez-Baeza Y, Pirrung M, González A & Knight R (2013) EMPeror: a tool for visualizing high-throughput microbial community data. GigaScience 2: 16. doi:10.1186/2047-217X-2-16

Vega FA, Covelo EF, Reigosa MJ, Andrade ML (2009) Degradation of fuel oil in saltmarsh soils affected by the Prestige oil spill. J Hazard Mater 166:1020–9 doi:10.1016/j.jhazmat.2008.11.113.

Vermuë M, Sikkema J, Verheul A, Bakker R & Tramper J (1993) Toxicity of homologous series of organic solvents for the gram-positive bacteria Arthrobacter and Nocardia sp. and the gram-negative bacteria Acinetobacter and Pseudomonas sp. Biotechnol Bioeng 42: 747–58. doi:10.1002/bit.260420610

Vieites DR, Nieto-Roman S, Palanca A, Ferrer X, Vences M (2004) European Atlantic: the hottest oil spill hotspot worldwide. Naturwissenschaften 91:535–8. doi:10.1007/s00114-004-0572-2

Vieira-Silva S, Rocha EPC (2010) The systemic imprint of growth and its uses in ecological (meta)genomic3.s. PLoSs. PLoS Genet 6(1):e1000808. doi:10.1371/ journal.pgen.1000808

Walker JD & Colwell RR (1975) Some effects of petroleum on estuarine and marine microorganisms. Can J Microbiol 21: 305–13.

Wang W & Shao Z (2013) Enzymes and genes involved in aerobic alkane degradation. Front Microbiol 4: 116. doi:10.3389/fmicb.2013.00116

Wang L, Wang W, Lai Q & Shao Z (2010a) Gene diversity of CYP153A and alkB alkane hydroxylases in oil-degrading bacteria isolated from the Atlantic Ocean. Environ Microbiol 12: 1230–42. doi:10.1111/j.1462-2920.2010.02165.x.

Wang W, Wang L & Shao Z (2010b) Diversity and abundance of oil-degrading bacteria and alkane hydroxylase (alkB) genes in the subtropical seawater of Xiamen Island. Microbial Ecol 60: 429–39. doi:10.1007/s00248-010-9724-4

Watts AW, Ballestero TP & Gardner KH (2006) Uptake of polycyclic aromatic hydrocarbons (PAHs) in salt marsh plants Spartina alterniflora grown in contaminated sediments. Chemosphere 62: 1253–60. doi:10.1016/j.chemosphere.2005.07.006

Wentzel A, Ellingsen T, Kotlar H-K, Zotchev S & Throne-Holst M (2007) Bacterial metabolism of long-chain n-alkanes. Appl Microbiol Biotech 76: 1209–21. doi:10.1007/s00253-007-1119-1

Whyte LG, Hawari J, Zhou E, Bourbonnière L, Inniss WE & Greer CW (1998) Biodegradation of variable-chain-length alkanes at low temperatures by a psychrotrophic Rhodococcus sp. Appl Environ Microbiol 64: 2578–84.

Whyte LG, Smits THM, Labbé D, Witholt B, Greer CW & van Beilen JB (2002) Gene cloning and characterization of multiple alkane hydroxylase systems in Rhodococcus strains Q15 and NRRL B-16531. Appl Environ Microbiol 68: 5933–42. doi:10.1128/AEM.68.12.5933-5942.2002

Wigand C, McKinney RA, Chintala MM, Charpentier MA & Groffman PM (2004) Denitrification Enzyme Activity of Fringe Salt Marshes in New England (USA). J. of Envt. Qual. 33(3): p 1144–1151. doi:10.2134/jeq2004.1144

Xu JL, He J, Wang ZC, Wang K, Li WJ, Tang SK & Li SP (2007) Rhodococcus qingshengii sp. nov., a carbendazim-degrading bacterium. Int J Syst Evol Microbiol 57: 2754–7. doi:10.1099/ijs.0.65095-0

Yan Y, Kuramae EE, Klinkhamer PGL, van Veen JA. (2015). Revisiting the dilution procedure used to manipulate microbial biodiversity in terrestrial systems. Appl Environ Microbiol 81: 4246–4252. doi:10.1128/AEM.00958-15

Zhu X, Venosa AD, Suidan MT & Lee K (2004) Guidelines for the Bioremediation of Oil-Contaminated Salt Marshes. Cincinnati, OH, US Environmental Protection Agency National Risk Management Research Laboratory Office of Research and Development. pp. 61

